# Managing river fish biodiversity generates substantial economic benefits in four European countries

**DOI:** 10.1101/447300

**Authors:** Carsten Riepe, Jürgen Meyerhoff, Marie Fujitani, Marie Fujitani, Øystein Aas, Johannes Radinger, Sophia Kochalski, Robert Arlinghaus

**Affiliations:** Department of Biology and Ecology of Fishes, Leibniz-Institute of Freshwater Ecology and Inland Fisheries, Müggelseedamm 310, D-12587 Berlin, Germany; Institute for Landscape Architecture and Environmental Planning, Technische Universität Berlin, Strasse des 17. Juni 145, D-10623 Berlin, Germany; Institutional and Behavioral Economics Working Group, Leibniz-Centre for Tropical Marine Research (ZMT), Fahrenheitstraße 6, D-28359 Bremen, Germany; Norwegian Institute for Nature Research, Fakkelgarden, N-2624 Lillehammer, Norway; GRECO, Institute of Aquatic Ecology, University of Girona, M. Aurèlia Capmany 69, SP-17003 Girona, Spain; Division of Integrative Fisheries Management, Albrecht-Daniel-Thaer-Institute of Agriculture and Horticulture & Integrative Research Institute for the Transformation of Human-Environment Systems, Faculty of Life Sciences, Humboldt-Universität zu Berlin, Invalidenstrasse 42, D-10115 Berlin, Germany

**Keywords:** choice experiment, economic values, river basin management plan, river fish conservation, native biodiversity, hydropower dams

## Abstract

Ecosystems and biodiversity produce benefits to society, but many of them are hard to quantify. For example, it is unclear whether European societies gain benefits from experiencing rivers that host high native biodiversity. Without such knowledge, monetary investments into ecologically oriented river management plans are difficult to justify. The objective of this study was to reveal how the public in four European countries values ecological characteristics of domestic rivers and the outcomes of hypothetical river basin management plans designed to improve river ecosystems, particularly fish biodiversity. We conducted a choice experiment among the populations in Norway, Sweden, Germany and France. We found similar preference structures in all countries with high marginal willingness-to-pay for improvements of abiotic river attributes (increased accessiblity of the river banks, improved bathing water quality, decreased river fragmentation). Citizens also benefited from certain fish species occurring in a river with native salmonid species being more valued than nonnatives, particularly in Norway, and from the degree of a river’s native biodiversity. Welfare measures calculated for selected river basin management plans (policy scenarios) revealed societal benefits that were primarily derived from ecological river management whereas a scenario focusing on hydroelectricity production generated the lowest utility. We conclude that ecological river management may produce high nonmarket economic benefits in all study countries, particularly through the management of abiotic river attributes and the restoration of declining or extinct fish species. Our results help to inform decisions on restoration efforts by showcasing the benefits that these measures have for the public.

## Introduction

Rivers and their fish populations deliver a range of ecosystem services (Holmlund and Hammer 1999; Auerbach et al. 2014), thereby contributing to human health and well-being (White et al. 2010; Nichols 2014). Due to a range of anthropogenic pressures (e.g., water abstraction, pollution, eutrophication, habitat degradation, damming, introduction of invasive species), the ecological status of many European watersheds, including the distribution of native fish species, has strongly declined over the last centuries (Dudgeon et al. 2006; Lenders et al. 2016). Today, riverine biodiversity has become one of the most threatened components of global biodiversity (Dudgeon et al. 2006; Collen et al. 2014), and ongoing economic development is further threatening river biodiversity in biodiversity hotspots (Zarfl et al. 2015; Winemiller et al. 2016). European freshwater fishes rank particularly high on the threat list relative to other vertebrates (Freyhof and Brooks 2011). Stressed ecosystems where biodiversity is in peril have been suggested to not deliver the full range of ecosystem services to society (Rockström et al. 2009; Sandifer et al. 2015), yet it is unclear to what extent this relationship applies to selected parts of the biotic world such as river fishes.

Ecological management ranks high on the priority list of many countries, which is reflected in their national policies and regulations aimed at curtailing biodiversity loss and restoring anthropogenically degraded ecosystems. In the European Union, the European Water Framework Directive (WFD; European Commission 2000), which has also been adopted by Norway, is an example of a regulation directed at the conservation of aquatic ecosystems. This policy aims at fostering the improvement of the ecological condition of aquatic ecosystems until a “good ecological status” is reached by 2027 (Hering et al. 2010; European Commission 2017). Recent assessments across Europe show that most surface waters fail to achieve a good status (or potential, for heavily modified or artificial water bodies), with rivers being generally in a worse condition than lakes (European Environment Agency 2018). The same is true for specific components of river biodiversity. For example, many riverine fish populations have strongly declined, and selected iconic species, such as Atlantic salmon (*Salmo salar*) and European eel (*Anguilla anguilla*) are threatened, and European sturgeon (*Acipenser* spp.) is virtually extinct across Europe (WWF 2001; Hindar 2003; Freyhof and Brooks 2011; Wolter 2015).

Conservation and restoration of these species demands considerable investment of public funds into river restoration (Szałkiewicz et al. 2018) like fish-friendly management of hydropower production (Nieminen et al. 2016). Such investments can only be justified if the public receives significant economic benefits from rivers with a good ecological status and from the presence of selected fish species. Estimating these benefits for the population at large requires an assessment of individual preferences for river development under trade-off conditions (European Commission 2000; Brouwer 2008). Given budgetary constraints, monetary values that the public associates with river attributes (e.g., the degree of biodiversity) can facilitate policy decisions on river basin management to meet ecological targets and to ensure that the costs are not disproportionate compared to the benefits that these actions generate for society (Brouwer 2008; Polizzi et al. 2015). One approach to eliciting the preferences of individual citizens for the status and future development of aquatic ecosystems are choice experiments (CE). CE are particularly well-suited for studying the trade-offs that precede preference formation in light of monetary constraints (Hanley et al. 2006; Brouwer 2008; Kataria 2009; Meyerhoff et al. 2014). We used a CE to evaluate the preferences for river attributes in four European countries (Norway, Sweden, Germany, France) and to understand whether ecological restoration goals align with nonmarket values attached by the general public to ecological river attributes as public goods. Modeling results were used to quantify the societal benefits of various policy scenarios as hypothetical outcomes of different river management strategies.

We assumed that people in all study countries prefer good water quality, easy access to the river banks and a high share of native biodiversity. We further expected that Scandinavians, particularly Norwegians, assign more value to specific fish species such as Atlantic salmon than the Germans or French because this species is economically and culturally more important in Norway, where it provides lucrative inland fisheries and aquaculture operations (WWF 2001; Hindar 2003) and has been receiving long-term coverage by mass media (Liu et al. 2016). By contrast, because species like Atlantic salmon are extinct in central Europe (e.g., in Germany) or strongly declining (as in France), central European citizens may have undergone an “extinction of experience” (Soga and Gaston 2016) and in turn may no longer benefit from knowing that a river is hosting salmon or other species they are hardly aware of (Kochalski et al. 2018).

## Materials and methods

### Choice experiment and survey instrument

A CE is a stated preference nonmarket valuation instrument that is consistent with utility maximization theory (Marschak 1960; McFadden 1974; Louviere et al. 2000) under the assumption that, given their budget constraints, people prefer one good over another good if the former maximizes the total expected utility gained from it. Due to the trade-offs implicit in a CE, the approach reveals more about respondents’ preferences and their underlying utility structures than asking directly and separately for preferences for individual attributes of a good, because people tend to want the best of everything (e.g., Daigle et al. 2016). CE are especially suitable for assessing public preferences for nonmarket goods and intrinsic values like those associated with biodiversity and other river attributes.

In our CE, we defined rivers as “running waters that are wide enough to allow for boating with small pleasure boats such as kayaks, canoes, or rowing boats”. Respondents were then presented with descriptions of two hypothetical river development programs (Fig. 1) that were specified along the levels of seven attributes (Table 1). To put the programs into temporal and spatial context, respondents were told that they would take effect within 10 years (Ahtiainen et al. 2015) and affect most rivers within a 50-km radius around their homes (Fig. 1). In the CE, we prompted respondents to consider only rivers within this area. A reference area defined by socially meaningful criteria (“home turf”; Liebich et al. 2018) promised to be more relevant to respondents than referring them to regions described in more biogeographical terms (like specific rivers or catchments; Liebich et al. 2018). Respondents were further informed that to achieve the outcome of a program, an obligatory financial contribution to a hypothetical river development fund was required, which the respondents would have to pay annually for a 10-year period (price attribute; Table 1; Fig. 1). Given the cost of each development program, respondents had to decide which one they preferred (Option A vs. B; Fig. 1), or alternatively, whether they wanted to maintain the current ecological status of the rivers (status quo) within their reference areas without any additional costs (Option C; Fig. 1). Respondents were asked about their subjective perception of the ecological status of these rivers before the CE was administered. The choice task thus required the respondents to trade off the total utility derived from one development program against that of the other program, depending on how much, if at all, they valued each of the river attribute levels. The estimated disutility of income loss (i.e., the price to be paid for a program) was used to rescale the utilities derived for the nonmonetary attributes to monitary units as a common metric, which made utilities directly comparable (Hanemann 1984). Prior to the presentation of the choice sets, respondents were introduced to all attributes (Table 2) and their levels (Table 1).

**Fig. 1.**
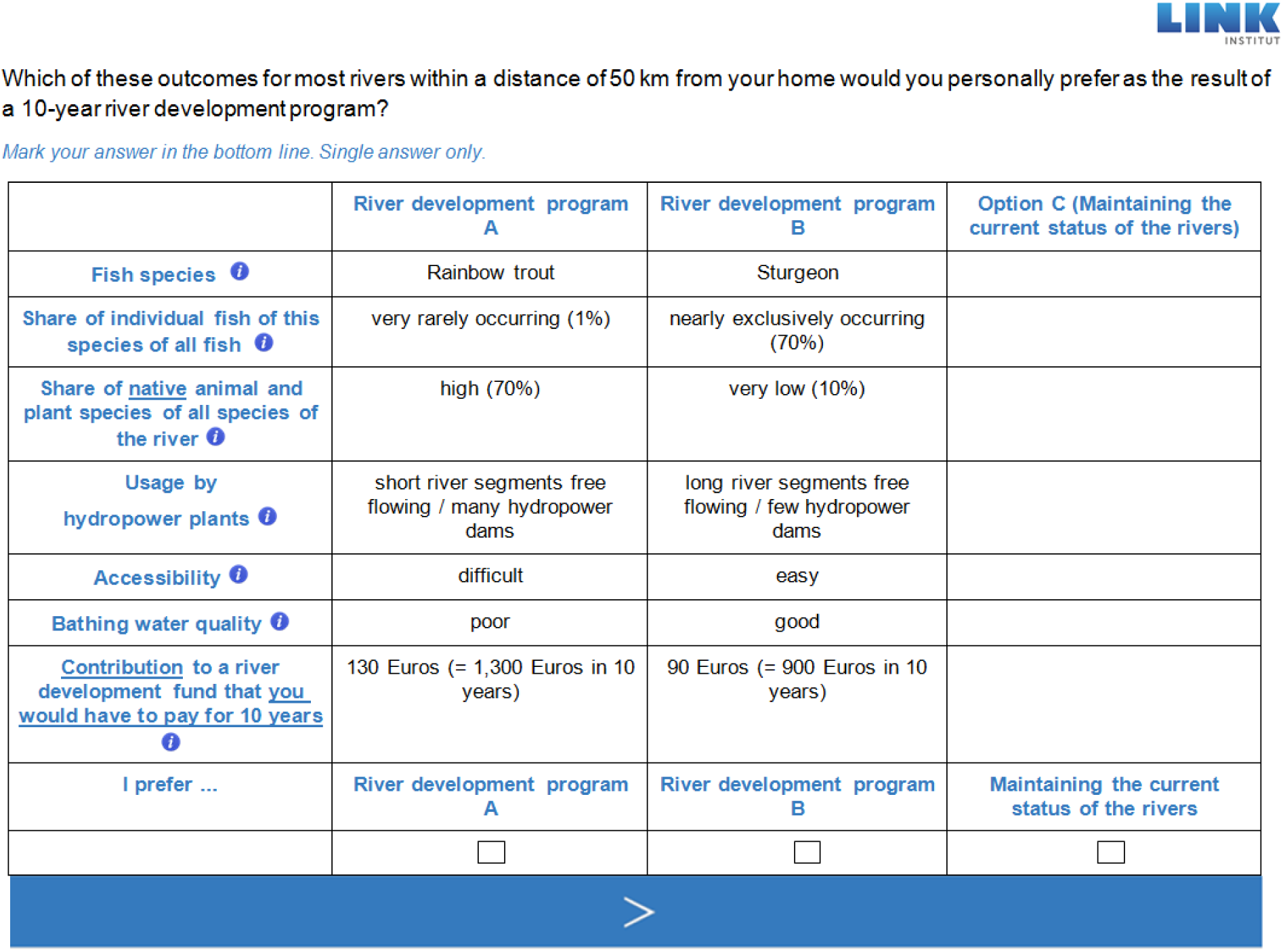
Example of a choice set as shown to respondents.

**Table 1.**
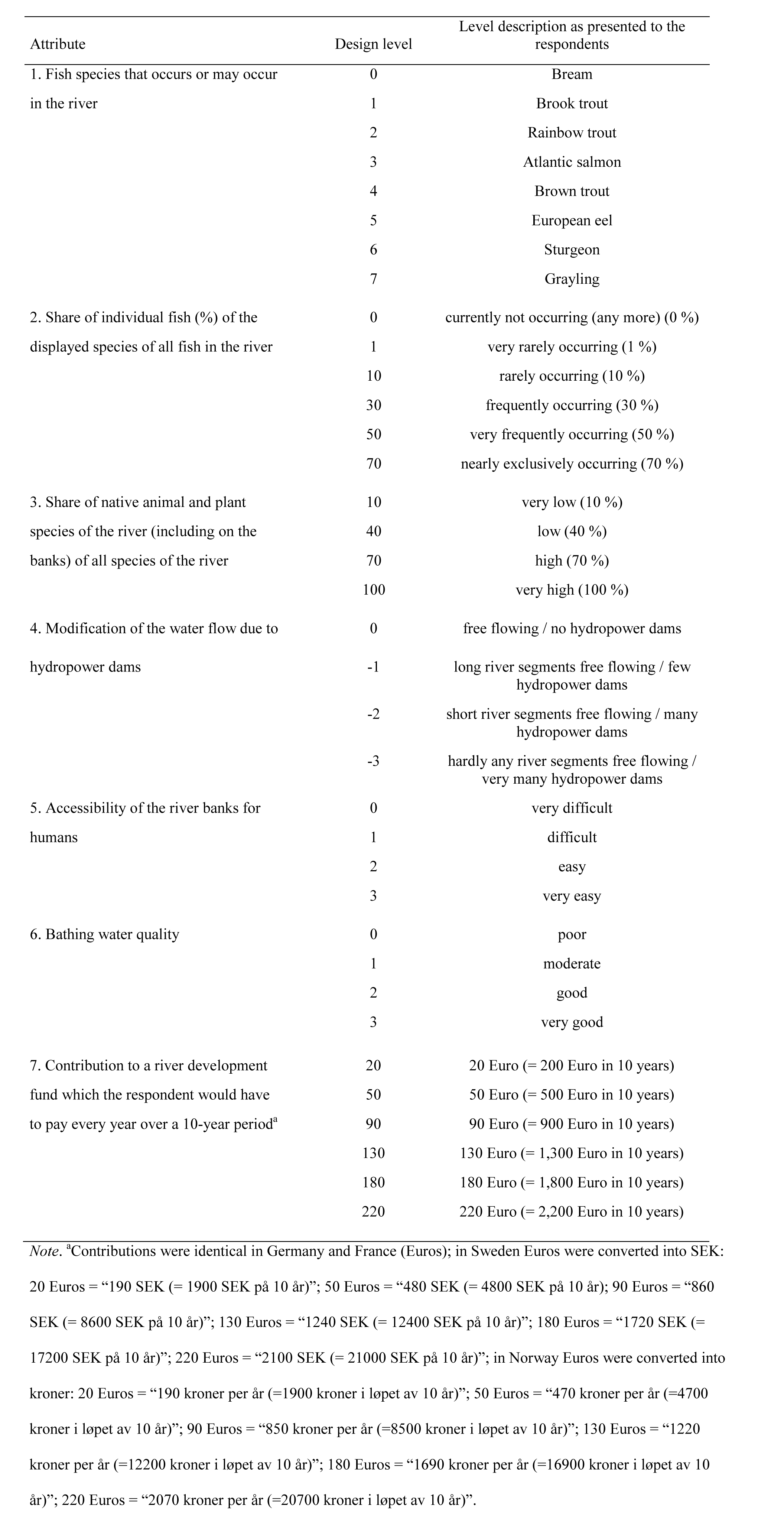
Attributes and design levels used in the choice experiment.

**Table 2.** Names of the attributes used in the choice experiment and introductory explanations as presented to the respondents.

| Short attribute name<sup>a</sup> | Long attribute name<sup>b</sup> | Attribute explanation given in the introduction of the respondents to the DCE<sup>c</sup> |
| --- | --- | --- |
| 1. Fish species | 1. Fish species that occurs or may occur in the river | Shows one of the fish species that, among others, occurs or may occur in the river. Some of the species are currently extinct and could be reintroduced through the release of hatchery-bred fishes (stocking). |
| 2. Share of individual fish of this species of all fish | 2. Share of individual fish (%) of the displayed species of all fish in the river | This is the relative frequency of the number of individuals of the fish species shown in relation to all individuals of all fish species in the river. A share of 0 % indicates that the species shown currently does not occur in the river (any more) and that it cannot be re-established given the river conditions shown. |
| 3. Share of native animal and plant species of all species of the river | 3. Share of native animal and plant species of the river (including on the banks) of all species of the river | This is the percentage of animals and plant species, including fish species, that are native to the river and its banks in relation to all species in the river. Native species have naturally colonized the river in the past without any human assistance. |
| 4. Usage by hydropower plants | 4. Modification of the water flow due to hydropower dams | Hydroelectric plants supply climate friendly electricity. They need dams which impound the water. Some fish species need free-flowing water to be able to migrate through a river to reach their spawning grounds. Dams and other transversal structures hamper these migrations or may even block them entirely, which may lead to species extinction. |
| 5. Accessibility | 5. Accessibility of the river banks for humans | The easier the access, the more of the river's shoreline can be walked on. River banks providing easy access for humans can result in the loss of natural habitats for plants and animals due to artificial embankments or effects of trampling. |
| 6. Bathing water quality | 6. Bathing water quality | Poor: turbid water, in summer occasional large area algal blooms, not suitable for swimming<br>Moderate: slightly turbid water, in summer occasional algal blooms, limited suitability for swimming<br>Good: largely clear water, suitable for swimming<br>Very good: very clear water, very suitable for swimming |
| 7. Contribution to a river development fund that you would have to pay for 10 years | 7. Contribution to a river development fund which you would have to pay every year over a 10-year period ... | ... to achieve the described river status for most rivers within 50 km from your residence. Note that the money that you would have to pay for a river development program would not be available for other expenditures any more. |

<sup>a</sup>Displayed on the choice sets (Fig. 1). <sup>b</sup>Used to introduce respondents to the choice experiment. <sup>c</sup>Explanations made available on each choice set via the info button next to the attribute's short name (Fig. 1).

The CE was constructed in a multi-stage development and pretest phase aiming to identify river attributes that indicate a good ecological status while being relevant to citizens’ everyday life. The attributes also had to be independent of each other and changeable through (hypothetical) management measures, thus allowing for policy scenario analyses. This phase involved experts from the study countries as well as exploratory interviews with members of the general population. The first of the final attributes that fulfilled the relevance criteria was the fish species occurring in a river (Attribute 1; Table 1). The levels of this attribute comprised two salmonid species (migratory Atlantic salmon and brown trout [*Salmo trutta*]; Table 1) that are native to all study countries. They can be considered flagship and umbrella species, that is, they are relevant to both fisheries and the general public, and indicative of a good ecological river status (Hindar 2003; Kalinkat et al. 2017). Economically important to Norway but extinct in Germany and endangered in France, Atlantic salmon is being restored in the latter two countries. It is also a familiar food product (WWF 2001; Hindar 2003; Wolter 2015), which nutritionists recommend for regular consumption (Dinter et al. 2016). We further included two nonnative salmonids (brook trout [*Salvelinus fontinalis*] and rainbow trout [*Oncorhynchus mykiss*]; Table 1) which were both introduced to Europe in the late 19th century (Aas et al. in press). These species are also known to the public as edible fishes but are often perceived as being native to Europe (Kochalski et al. 2018). Two further migratory fish species were included: threatened European eel and the extinct sturgeon (Table 1), which benefit from free flowing rivers when migrating to their spawning grounds (Nieminen et al. 2016). Grayling (*Thymallus thymallus),* another native salmonid, and bream (*Abramis brama),* a cyprind, were also included (Table 1). These species were not expected to strongly increase river utility, but they are key species determining fish regions in European rivers (Huet 1949). Another biological attribute was the relative abundance of each species shown on the choice sets (defined as the “share of individual fish”; Attribute 2; Table 1). This attribute’s levels ranged from 0% to 70% to include even very high levels of abundance. We also included native biodiversity as a more generic attribute (“share of native animal and plant species"; Attribute 3; Table 1), whose levels ranged from 10% to 100%, assuming that its highest level was ecologically most valuable. Referring directly to the presence of riverine organisms, these three attributes reflected biotic river characteristics.

The remaining four attributes described abiotic river characteristics, primarily reflecting the human perspective on the use of rivers while still being closely related to biological river conditions. We included the degree of modification of the water flow due to hydropower dams as fourth attribute (Table 1), as it threatens the natural ecological function of rivers across the world (Zarfl et al. 2015; Winemiller et al. 2016; Couto and Olden 2018) and particularly migratory fishes (Lawrence et al. 2016; Cooper et al. 2017). This attribute implicitly required respondents to compare the utility derived from the production of climate-friendly electricity with the utility gained from knowing that fishes were able to migrate (Table 2) and other (e.g., aesthetic) values potentially associated with a free-flowing river. A free-flowing river was taken to be ecologically most valuable. Accessibility of the river banks (Attribute 5; Table 1) and the bathing water quality (Attribute 6; Table 1) were also deemed to be related to a good river status as perceived by the public (Hanley et al. 2006; Kataria 2009; Artell and Huhtala 2017). The definitions of the attribute levels of bathing water quality (Table 2) were adapted from the water quality ladder used by Meyerhoff et al. (2014). Respondents were instructed to consider this attribute to be independent of whether they actually used a river for bathing or not. Riparian zones of rivers provide natural habitats for plants and animals, which can be destroyed through artificial embankments or effects of trampling (Arlinghaus et al. 2002; Tockner et al. 2010). Very difficult access to the river banks and very good bathing water quality were thus thought to indicate a river’s good ecological status. The obligatory annual contribution to a hypothetical river development fund served as price attribute (Attribute 7; Table 1).

To familiarize respondents with the attributes and their levels, we ascertained their perceptions of the status quo of the rivers within their reference areas using rating scales with verbal descriptors identical to the attribute levels used in the CE (Ahtiainen et al. 2015). This was done for the share of native animal and plant species of the river, the modification of the water flow due to hydropower plants, the accessibility of the banks, and the bathing water quality (Attributes 3 to 6; Table 1). As we expected most study participants to have only little, if any, knowledge of the fish species assemblage in their nearby rivers (Kochalski et al. 2018; Liebich et al. 2018), we did not ask for the assumed presence of particular fish species and their abundance (Attributes 1 and 2; Table 1).

Bayesian efficient statistical designs were created for a multinomial logit model (Scarpa and Rose 2008) to allocate attribute levels (Table 1) to river development programs to fully enumerate respondents’ preferences for different attributes. As design criteria, we used D-efficiency and S-efficiency for which we created 32 choice sets each that were blocked into four distinct subsets each encompassing eight choice sets. Two design criteria were used to mitigate potential biases due to optimizing only for one criterion (Olsen and Meyerhoff 2016). Respondents were randomly assigned to one block of choice sets. The questionnaire also mapped the demographic background of the respondents.

### Data analysis

The analysis of the stated choices is based on the random utility model (McFadden 1974). It assumes that an individual decision maker’s preferences are the sum of a systematic (*V*) and an unobservable or stochastic component (*ε*), where *V* is an indirect utility function. If the stochastic component is distributed independently and identically and follows a Gumbel distribution, the conditional probability that alternative *i* is chosen by individual *n* is defined as:

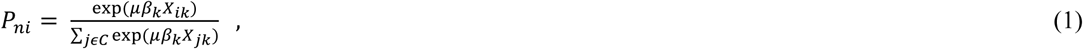

where the scale parameter (*μ*) of the error distribution is confounded with the parameter vector *β_k_* and generally normalized to 1, *X_ik_* is attribute *k* of alternative *i*. As the simple logit is unable to account for unobserved taste heterogeneity and the fact that each respondent faced 8 choice sets (leading to repeated measures), we opted for a mixed logit model. This model is an extension of the basic multinomial logit model estimating not only the mean for each attribute parameter but also the deviation of each respondent from the sample mean taking unobserved taste heterogeneity into account (Train 2009). For all nonmonetary attributes, we assumed that the parameters specified as random follow a normal distribution. The cost attribute, however, was set to follow a lognormal distribution as it ensures that the coefficient has always the same sign; the cost attribute was multiplied by −1 before estimation. We also investigated observed taste heterogeneity and included interactions between the alternative-specific constant for the current situation (ASCsq) and respondent-related characteristics. These comprised sociodemographic (age, gender, education) as well as environmental characteristics (land use and prevalence of rivers within the 50-km reference areas). Significant coefficients for the interactions indicate an influence of these characteristics on the likelihood that the status quo of the rivers (Option C; Fig. 1) is chosen instead of a program that would change the current ecological conditions in the river. To determine the environmental characteristics for each respondent individually, we sourced geographical information about the degree of urbanization and the number of rivers within their 50-km areas using GRASS GIS (Neteler et al. 2012). We extracted geographical coordinates for the zipcode of each respondent and obtained land cover information from the European Corine Land Cover (CLC) 2012 database at a spatial resolution of 250×250m (http://land.copernicus.eu/pan-european/corine-land-cover/clc-2012/view). We also obtained a vector river network from the European CCM river and catchment database (http://ccm.jrc.ec.europa.eu; de Jager & Vogt 2010). For further analysis the original CLC classes 1 to 11 were aggregated to a single thematic class representing urbanization. Subsequently, we calculated (i) the percentage of urban land cover and (ii) the number of unique rivers within a buffer radius of 50 km around each respondent’s location. These data were matched with the survey data.

As the status quo of the rivers is expected to vary between respondents’ reference areas due to natural causes, the assumption of a uniform status quo for all respondents can lead to biased coefficient and welfare estimates. We therefore constructed individualized status-quo alternatives during the data modeling process according to the individual perception of the current river conditions as reported by each respondent. Because we used the same attribute levels for the description of the two river development programs on the choice sets (Options A and B; Fig. 1; Table 1) and the questionnaire-based assessment of the status quo, we were able to use distinct attribute levels to define each respondent’s status-quo alternative (Option C; Fig. 1). If status-quo data were missing, we imputed them countrywise with the means of all respondents with nonmissing data (Ahtiainen et al. 2015). All model parameters were estimated by simulated maximum likelihood using Halton draws with 1000 replications.

Conversion of parameter estimates into marginal willingness-to-pay (MWTP) facilitates the comparison of parameters between countries because a common monetary scale unit (€) is used for all attributes. We estimated the MWTP values as the negative ratios of the attribute parameters and the cost parameter (Hanemann 1984). They indicate how much respondents were willing to pay for a one-level change of a nonmonetary attribute, for example, from moderate to good bathing water quality quantifying the desirability of perceived benefits from the level change (Table 1). To calculate the MWTP estimates, we used the median value of the log-normally distributed price coefficient because using its mean value would have resulted in unreasonably low MWTP estimates close to zero. A model with a fixed price coefficient, in turn, would result in significantly lower model fit as it assumes that no heterogeneity exists among respondents towards cost. The median, on the other hand, is more robust to extreme values (Bliemer and Rose 2013), and the estimated coefficients are consistent with the estimated price coefficient of the mixed logit model with a fixed price coefficient (Sagebiel et al. 2017).

Subsequently, we calculated nonmarginal welfare measures (Hanemann 1984) for a range of policy scenarios. The measures indicate the benefits accrued to society from a given combination of attribute level changes relative to the status quo:

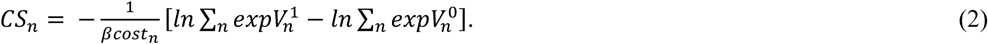

Here, *CS* is the compensating surplus welfare measure, *β_cost_* is the marginal utility of income (the coefficient of the cost attribute) and *V^0^_n_* and *V^1^ _n_* represent the *n*th individual’s indirect utility functions before and after the change under consideration. We used the 95% confidence intervals to determine the statistical significance of all MWTP differences, and of within-country CS differences. Between-country CS differences were tested for significance using the Poe test (Poe et al. 2005).

### Policy scenarios

We developed six policy scenarios to understand the population benefits in terms of CS values that may result from the ecological outcomes of distinct 10-year river basin management plans (Table 3). The resulting CS values reflect the joint effect of each scenario’s combination of attribute levels relative to the baseline levels (for Attributes 1 and 2; Table 1) and to the individual status-quo levels (as ascertained for Attributes 3 to 6; Table 1), respectively, that were assumed for Option C (Fig. 1). The scenarios were set up according to different management strategies focusing either on (i) improved conditions for capture fisheries (Scenarios 1 and 2), (ii) nature conservation (Scenarios 3 and 4), or (iii) green-energy production through hydropower plants (Scenarios 5 and 6; Table 3), alongside assumed impacts on fishes and general river biodiversity. By comparing the utilities of possible outcomes, our analyses showcase the benefits that river restoration may bring about in each of the four countries.

**Table 3.** Description of policy scenarios for six alternative river basin management plans.

| Managerial orientation |  |  |  |  |  |  |
| --- | --- | --- | --- | --- | --- | --- |
|  | Scenario 1<br>Fisheries<br>focusing on<br>native<br>salmonids | Scenario 2<br>Fisheries<br>focusing on<br>nonnative<br>salmonids | Scenario 3<br>Conservation-oriented<br>focusing on<br>native<br>salmonids | Scenario 4<br>Focusing on<br>holistic ecosystem<br>conservation | Scenario 5<br>Hydropower<br>(Green Energy) | Scenario 6<br>Hydropower (Green<br>Energy) and<br>fisheries focusing on<br>native salmonids |
| Attribute | Attribute levels |  |  |  |  |  |
| 1. Fish species | Atlantic salmon | Rainbow trout | Atlantic salmon | Atlantic salmon | Bream | Brown trout |
| 2. Share of individual fish of<br>the displayed species of all<br>fish | very frequently<br>occurring (50<br>%) | frequently<br>occurring (30<br>%) | frequently<br>occurring (30<br>%) | rarely occurring<br>(10 %) | very frequently<br>occurring (50 %) | frequently occurring<br>(30 %) |
| 3. Share of native animal and<br>plant species of all species of<br>the river | high (70 %) | low (40 %) | very high (100<br>%) | very high (100 %) | very low (10 %) | high (70 %) |
| 4. Usage by hydropower<br>plants | long river<br>segments free<br>flowing / few<br>hydropower<br>dams | short river<br>segments free<br>flowing / many<br>hydropower<br>dams | long river<br>segments free<br>flowing / few<br>hydropower<br>dams | free flowing / no<br>hydropower dams | hardly any river<br>segments free<br>flowing / very<br>many hydropower<br>dams | short river segments<br>free flowing / many<br>hydropower dams |
| 5. Accessibility | easy | very easy | difficult | very difficult | very easy | easy |
| 6. Bathing water quality | good | moderate | good | very good | moderate | very good |

Both fisheries-oriented scenarios (Scenarios 1 and 2; Table 3) focused on salmonid fish species to maintain capture fisheries. The native-salmonid scenario (Scenario 1) had Atlantic salmon occuring very frequently, which would also benefit other riverine species. Therefore, this scenario also assumed a high level of native biodiversity and medium levels of the three abiotic attributes (Attributes 4 to 6; Table 3). The nonnative-salmonid fisheries scenario (Scenario 2) featured rainbow trout, which hardly reproduce in central Europe and therefore need to be stocked. This scenario assumed a correspondingly low share of native biodiversity, very easy accessibility of the banks for fishers to be able to reap the benefits of rainbow trout stocking and medium levels of the other two attributes (Table 3). The conservation-oriented scenarios (Scenarios 3 and 4; Table 3) were unrelated to fisheries. They included migratory Atlantic salmon as a flagship umbrella species (Kalinkat et al. 2017), indicating a good ecological river status, and had very high levels of native biodiversity (Table 3). In the native-salmonids conservation scenario (Scenario 3), Atlantic salmon occurred frequently in rivers with only few hydropower dams and good bathing water quality to render this scenario comparable to the native fisheries scenario (Scenario 1), but here we assumed difficult access to the river banks to improve the ecological quality of the riparian zone (Table 3). In the holistic-ecosystem scenario (Scenario 4), we assumed Atlantic salmon to occur less frequently, and for the three abiotic attributes we assumed the levels that we considered ecologically most valuable (i.e., no hydropower dams, very difficult accessibility, very good water quality; Table 3). The green-energy scenarios (Scenarios 5 and 6; Table 3) focused on the production of climate-friendly electricity from hydropower plants. Consequently, hydropower Scenario 5 assumed very many hydropower dams. As no emphasis was put on fisheries in this scenario, we included frequently occurring bream as fish species, which is not often targeted by fishers and whose abundance increases in flow-regulated rivers, along with a very low level of native biodiversity, very easy accessibility and moderate water quality (Table 3). In Scenario 6, we maintained the goal of green-energy production while facilitating capture fisheries through regulated hydropower, comparable to the situation in Norway (Alfredsen et al. 2012). To that end, we assumed a reduced number of hydropower dams and ecologically improved levels of the other two abiotic attributes compared to hydropower Scenario 5. In addition, we assumed native brown trout, another flagship and umbrella species often targeted by fishers and indicative of a good ecological river status, to occur frequently together with a high level of native biodiversity (Table 3).

### Sample and data collection

The questionnaire was administered by means of an internet-based survey that was carried out in September 2015 among the general populations aged 16 to 74 years living in private households in Norway, Sweden, Germany and France (*n*=1,000 per country). Study participants were randomly sampled from online consumer panels (with 40,000 to nearly 100,000 members per country) whose members had been previously recruited by phone (i.e., offline) using a probability-based, random digit-dialing method as sampling frame (Heckel et al. 2014; ADM 2018). The online populations (i.e., persons living in households with internet access) covered between 83% (France) and 97% (Norway) of all private households (Germany: 90%; Sweden: 91%; Eurostat 2016). Country-specific quotas were set on age groups, gender and the highest education level achieved (as standardized by the International Classification of Education ISCED; UNESCO 2016) according to census data (Eurostat 2015). Fieldwork including the development and administration of the questionnaire was planned and conducted following recommendations given by Dillman et al. (2014). The data collection phase was preceded by technical pretests and *n*=30 pilot interviews per country. Participants of the main study were invited by email followed by up to three reminder emails.

## Results

### Sample description

The samples did not differ significantly in mean age (ranging from 41.5 years in France to 43.2 years in Sweden; post-hoc tests: *p* ≥ .05; *F* = 2.7, *df* = 3) and gender composition (Table 4) but slightly differed in education levels (Table 4). These distributions mirror the online populations of the four countries according to census data (Eurostat 2015). Norwegian and Swedish respondents perceived the status quo of their nearby rivers quite similarly. Both assumed the rivers’ native biodiversity to be higher, the accessibility of the banks easier and the bathing water quality better than the Germans and particularly the French (Table 5). While respondents in all countries characterized the water flow of the rivers as modified by only a few hydropower dams (Table 5), the Norwegians considered the water flow to be somewhat stronger modified than the Swedish and German respondents. The French perceived the least dam-related impact on the water flow (Table 5).

**Table 4.** Key characteristics of the samples (%).

| | France | Germany | Norway | Sweden | $\chi^2$ (df) |
| --- | --- | --- | --- | --- | --- |
| Gender |  |  |  |  | 0.5 (3) |
| female | 48.9 | 48.4 | 47.5 | 47.7 |  |
| male | 51.1 | 51.6 | 52.5 | 52.3 |  |
| Education level<sup>a</sup> |  |  |  |  | 94.4 (6) * |
| low (0 – 2) | 17.8 | 11.1 | 18.4 | 18.2 |  |
| medium (3 or 4) | 45.6 | 59.0 | 38.6 | 44.1 |  |
| high (5 – 8) | 36.6 | 29.9 | 43.0 | 37.7 |  |

<sup>a</sup>According to the International Classification of Education (ISCED; UNESCO, 2016).
\* $p < .05$ .

**Table 5.** Perceived status quo of the rivers within a 50-km area around respondents’ homes: means (*M*), standard deviations (*SD*) and ANOVA for river attributes used in the choice experiment.

| River attribute | France<br>min. *n* = 647<br>max. *n* = 831 |  | Germany<br>min. *n* = 853<br>max. *n* = 963 |  | Norway<br>min. *n* = 605<br>max. *n* = 845 |  | Sweden<br>min. *n* = 626<br>max. *n* = 873 |  | ANOVA |  |
| --- | --- | --- | --- | --- | --- | --- | --- | --- | --- | --- |
|  | *M* | *SD* | *M* | *SD* | *M* | *SD* | *M* | *SD* | *F* | df |
| Share of native animal and plant species of all species of the river<sup>a</sup> | 2.2<sub>a</sub> | 0.7 | 2.4<sub>b</sub> | 0.6 | 2.7<sub>c</sub> | 0.7 | 2.6<sub>c</sub> | 0.7 | 63.1 * | 3, 2726 |
| Usage by hydropower plants<sup>b</sup> | 1.7<sub>a</sub> | 0.8 | 1.9<sub>b</sub> | 0.6 | 2.1<sub>c</sub> | 0.9 | 1.9<sub>b</sub> | 0.8 | 33.9 * | 3, 2922 |
| Accessibility<sup>c</sup> | 2.8<sub>a</sub> | 0.7 | 2.9<sub>a</sub> | 0.7 | 3.2<sub>b</sub> | 0.6 | 3.2<sub>b</sub> | 0.7 | 84.7 * | 3, 3489 |
| Bathing water quality<sup>d</sup> | 1.9<sub>a</sub> | 0.8 | 2.4<sub>b</sub> | 0.7 | 2.6<sub>c</sub> | 0.9 | 2.6<sub>c</sub> | 0.8 | 131.3 * | 3, 3499 |

*Note.* Means in each row that share subscripts do not differ significantly ( $p \geq .05$ ; Games-Howell test, Hochberg's GT2 test).
<sup>a</sup>Scale from 1 (*very low* [10 %]) to 4 (*very high* [100 %]). <sup>b</sup>Scale from 1 (*free flowing / no hydropower dams*) to 4 (*hardly any river segments free flowing / very many* *hydropower dams*). <sup>c</sup>Scale from 1 (*very difficult*) to 4 (*very easy*). <sup>d</sup>Scale from 1 (*poor*) to 4 (*very good*). For descriptors of intermediate scale levels see Table 1.
\* $p < .05$ .

### Preferences for river attributes

The negative parameter estimates for the ASCsq in all countries (Table 6) indicated that the respondents derived utility from moving away from the status quo and thus from choosing to contribute financially to a river development program. Coefficients for the interactions between the ASCsq and the sociodemographic and environmental characteristics showed mixed results. While the utility of the status-quo alternative increased with increasing age in all countries, and with being female in France, it decreased for female respondents in Germany (Table 6). Furthermore, its utility decreased in Norway and Sweden with increasing degree of urbanization and increased in Norway with increasing number of nearby rivers (Table 6).

**Table 6.**
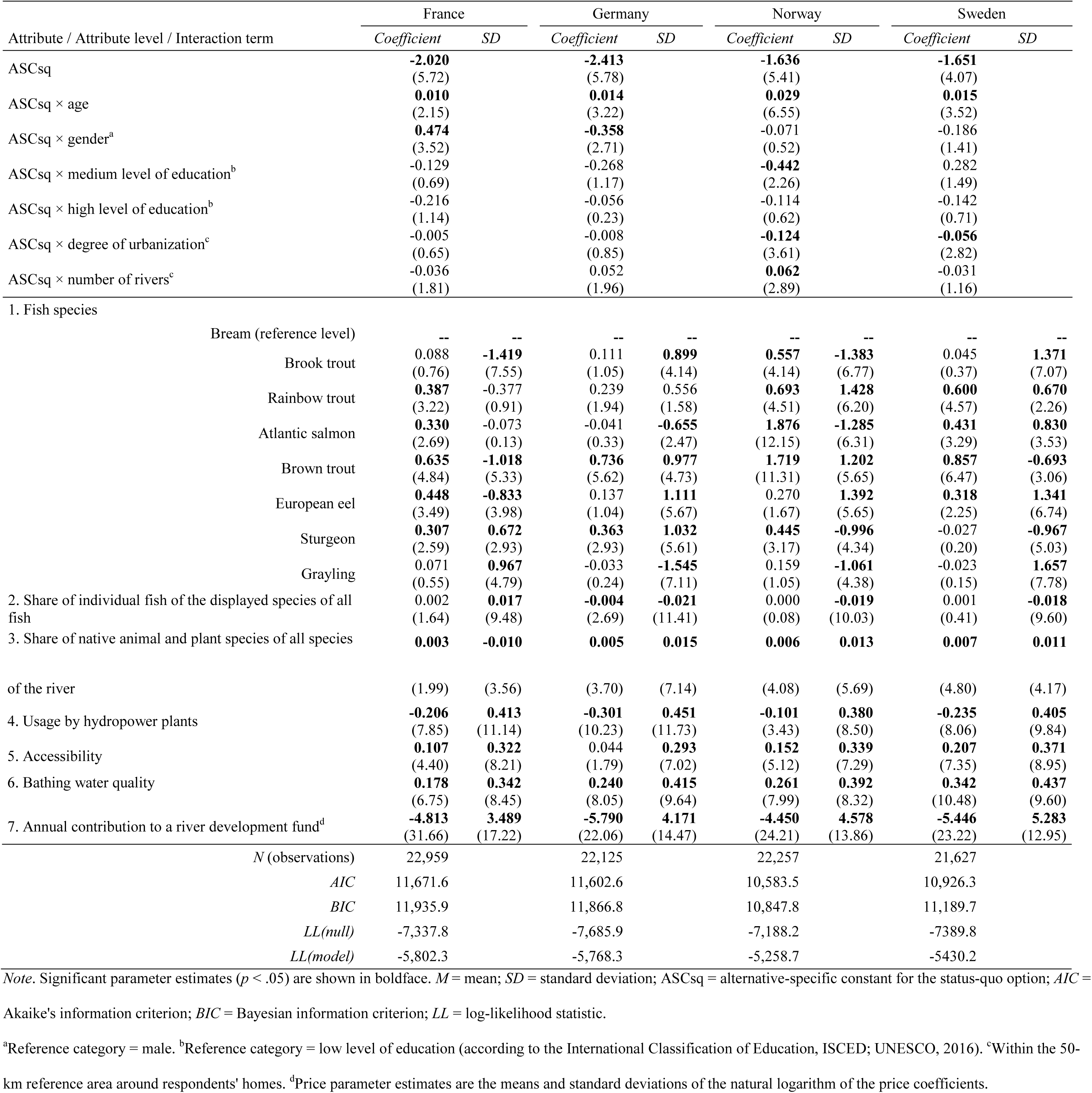
Estimated model coefficients and standard deviations (*SD*) for attributes and attribute levels used in the choice experiment (*z* values in parentheses).

Except for grayling, all fish species contributed to a river’s perceived utility, relative to bream, in at least one country. Five species provided benefits in France and in Norway, four in Sweden and two in Germany (Table 6). The native salmonid species (Atlantic salmon, brown trout) were generally more appreciated than the nonnatives (brook and rainbow trouts) as evidenced by the total number of significant parameters across all countries (seven vs. four, respectively; Table 6). Also, utility was derived from European eel in France and Sweden and by sturgeon in France, Germany and Norway (Table 6). The relative abundance of a fish species in a river did not impact utility, except in Germany where its influence was negative (Table 6). An increase in the share of native biodiversity and in bathing water quality increased utility in all countries as did an increase in accessibility of the river banks, except in Germany (Table 6). The more a river’s water flow was modified due to hydropower dams, the more the expected utility of a river decreased in all countries (Table 6).

In line with economic theory, the negative sign of the cost attribute indicates the decreasing probability of a respondent to choose an alternative when its price rises (Table 6). In all countries, the standard deviations for most attributes were significant, in some instances solely the standard deviation became significant (Table 6). As the random parameter model captures unobserved taste heterogeneity with respect to the attributes, significant standard deviations indicate the presence of taste heterogeneity in the sample. The model results thus bear witness to considerable unobserved taste heterogeneity among respondents implying strong differences in preferences within the populations.

### Marginal willingness-to-pay (MWTP) for river attributes

A comparison of the MWTP values within fish species (Attribute 1) across countries as well as across species within countries resulted in only few significant differences as evidenced by nonoverlapping confidence intervals (Table 7). Within-country differences were found only in Norway where native salmonids (Atlantic salmon: 160.7 €, brown trout: 147.2 € per year) were valued higher than sturgeon and nonnative brook and rainbow trouts (38.2 €, 47.7 € and 59.4 € per year, respectively). The Norwegian MWTP for Atlantic salmon was also the only species-related value that differed significantly from its corresponding value in another country (France: 40.6 € per year; Table 7). An increase in the relative abundance of the focal fish species (Attribute 2) affected a river’s utility only in Germany where it led to a decrease in MWTP by 1.3 € per year per *%* increase. An increase in native river biodiversity (Attribute 3) increased river utility in Germany, Norway and Sweden by 1.7, 0.5 and 1.5 € per year, respectively, per *%* increase (Table 7). As for the MWTP values of the abiotic attributes, which quantify the change in utility for a one-level increment of an attribute, countries differed strongly in the disutility entailed by an increase in the number of hydropower dams (Attribute 4). Whereas a one-level increase decreased the amount of money people would be willing to pay for a river development plan in Germany by 98.3 € per year, the MWTP in Norway decreased by only 8.6 € per year (Table 7). The negative utility of this attribute in France was higher than the latter (25.4 € per year), while the Swedish MWTP decreased by 54.5 € per year (Table 7). MWTP values associated with a one-level increase in the accessibility of the river banks (Attribute 5) and in bathing water quality (Attribute 6) increased by approximately the same amounts in both France and Norway (accessibility: 13 € per year; bathing water quality: 22 € per year; Table 7). For bathing water quality, economic values in Germany and Sweden were higher and also very similar (79 € per year; Table 7). The MWTP value of the accessibility in Sweden was 48 € per year, while this attribute did not significantly add to a river’s perceived utility in Germany (Table 7).

**Table 7.** Marginal willingness-to-pay (MWTP) estimates (€ / year) by country for attributes and attribute levels.

| Attribute / Attribute level | France |  |  | Germany |  |  | Norway<sup>a</sup> |  |  | Sweden<sup>a</sup> |  |  |
| --- | --- | --- | --- | --- | --- | --- | --- | --- | --- | --- | --- | --- |
|  | MWTP | 95% CI |  | MWTP | 95% CI |  | MWTP | 95% CI |  | MWTP | 95% CI |  |
|  |  | lower limit | upper limit |  | lower limit | upper limit |  | lower limit | upper limit |  | lower limit | upper limit |
| 1. Fish species |  |  |  |  |  |  |  |  |  |  |  |  |
| Brook trout | 10.8 | -17.2 | 38.8 | 36.2 | -33.2 | 105.7 | **47.7\*** | 20.3 | 75.1 | 10.4 | -45.4 | 66.2 |
| Rainbow trout | **47.6\*** | 15.9 | 79.3 | 78.3 | -9.0 | 165.5 | **59.4\*** | 27.0 | 91.8 | **139.0\*** | 54.7 | 223.2 |
| Atlantic salmon | **40.6\*** | 9.1 | 72.2 | -13.5 | -95.1 | 68.0 | **160.7\*** | 100.6 | 220.8 | **100.0\*** | 26.8 | 173.2 |
| Brown trout | **78.1\*** | 39.4 | 116.9 | **240.9\*** | 93.6 | 388.2 | **147.2\*** | 92.4 | 202.1 | **198.7\*** | 92.7 | 304.8 |
| European eel | **55.2\*** | 20.0 | 90.4 | 44.9 | -43.6 | 133.5 | 23.1 | -4.9 | 51.1 | **73.7\*** | 1.2 | 146.2 |
| Sturgeon | **37.7\*** | 6.9 | 68.6 | **118.9\*** | 16.5 | 221.4 | **38.2\*** | 11.4 | 65.0 | -6.2 | -65.7 | 53.3 |
| Grayling | 8.8 | -22.6 | 40.1 | -10.9 | -101.0 | 79.3 | 13.6 | -11.9 | 39.1 | -5.2 | -75.6 | 65.2 |
| Bream | - | - | - | - | - | - | - | - | - | - | - | - |
| 2. Share of individual fish of the displayed species of all fish<sup>b</sup> | 0.3 | -0.1 | 0.6 | **-1.3\*** | -2.4 | -0.1 | 0.0 | -0.2 | 0.3 | 0.1 | -0.5 | 0.8 |
| 3. Share of native animal and plant species of all species of the river<sup>b</sup> | 0.3 | -0.0 | 0.7 | **1.7\*** | 0.4 | 3.0 | **0.5\*** | 0.2 | 0.8 | **1.5\*** | 0.6 | 2.5 |
| 4. Usage by hydropower plants<sup>c</sup> | **-25.4\*** | -35.1 | -15.8 | **-98.3\*** | -151.9 | -44.8 | **-8.6\*** | -14.3 | -2.9 | **-54.5\*** | -82.7 | -26.3 |
| 5. Accessibility<sup>c</sup> | **13.1\*** | 6.1 | 20.1 | 14.3 | -3.1 | 31.7 | **13.0\*** | 6.4 | 19.6 | **47.9\*** | 22.8 | 73.1 |
| 6. Bathing water quality<sup>c</sup> | **21.9\*** | 12.9 | 30.9 | **78.5\*** | 33.6 | 123.4 | **22.4\*** | 12.5 | 32.2 | **79.3\*** | 40.0 | 118.6 |

*Note.* Significant MWTP estimates are shown in boldface. CI = confidence interval.
<sup>a</sup>MWTP values in NOK and SEK, respectively, were converted to € <sup>b</sup>Increase in MWTP per % increase. <sup>c</sup>Increase in MWTP for a one-level increase.
\* $p < .05$ .

### Benefits of the policy scenarios

All scenarios provided nonzero benefits to the four societies except for the hydropower (green-energy) Scenario 5 in Germany (Table 8). In the other three countries, this scenario still delivered the lowest CS values of all scenarios. Moreover, it was the only one whose CS values showed 95%-confidence intervals that did not overlap with those of other scenarios in the same country. Except in Sweden, the hydropower (green-energy) Scenario 5 delivered lower benefits than the scenario focusing on fisheries for native salmonids (Scenario 1) and both conservation-oriented scenarios (Scenarios 3 and 4; Table 8). Apart from the value-lowering presence of very many hydropower dams (Table 7), the low utilities derived from the hydropower (green-energy) scenario also originated from moderate bathing water quality, very low share of native biodiversity and very frequent occurrence of bream, the reference species (Table 3). The joint CS-diminishing effect of these attribute levels, however, was compensated for as soon as the green-energy management strategy underlying hydropower Scenario 5 was modified to also attain the goal of facilitating capture fisheries for native brown trout, alongside improvements in native biodiversity and bathing water quality (Scenario 6; Tables 3, 8). In consequence, nonoverlapping confidence intervals between both hydropower scenarios in all four countries indicated a significant increase in CS from a strict hydropower management strategy (Scenario 5) to a strategy that additionally facilitated fisheries (Scenario 6; Table 8).

**Table 8.** Compensating surplus (CS) in € per year for six policy scenarios.

| Scenario<sup>b</sup> | France |  |  | Germany |  |  | Norway<sup>a</sup> |  |  | Sweden<sup>a</sup> |  |  |
| --- | --- | --- | --- | --- | --- | --- | --- | --- | --- | --- | --- | --- |
|  | CS | 95% CI |  | CS | 95% CI |  | CS | 95% CI |  | CS | 95% CI |  |
|  |  | lower limit | upper limit |  | lower limit | upper limit |  | lower limit | upper limit |  | lower limit | upper limit |
| Scenario 1<br>Fisheries native salmonids | 197.5*<sub>a</sub> | 125.4 | 269.6 | 425.9*<sub>bc</sub> | 175.6 | 676.1 | 284.2*<sub>ab</sub> | 178.5 | 389.9 | 578.7*<sub>c</sub> | 303.0 | 854.4 |
| Scenario 2<br>Fisheries nonnative salmonids | 155.2*<sub>a</sub> | 96.8 | 213.7 | 328.8*<sub>bd</sub> | 130.3 | 527.4 | 149.7*<sub>ac</sub> | 88.4 | 211.0 | 482.6*<sub>d</sub> | 252.4 | 712.8 |
| Scenario 3<br>Conservation native salmonids | 189.0*<sub>a</sub> | 117.9 | 260.2 | 488.2*<sub>bc</sub> | 206.6 | 769.8 | 285.9*<sub>ab</sub> | 179.2 | 392.6 | 574.6*<sub>c</sub> | 299.8 | 849.4 |
| Scenario 4<br>Holistic Ecosystem Conservation | 218.1*<sub>a</sub> | 139.3 | 296.9 | 675.9*<sub>bd</sub> | 303.3 | 1048.6 | 303.7*<sub>ac</sub> | 190.3 | 417.0 | 657.8*<sub>d</sub> | 344.4 | 971.3 |
| Scenario 5<br>Hydropower (Green Energy) | 77.6*<sub>a</sub> | 43.4 | 111.8 | 75.6<sub>a</sub> | -6.6 | 157.8 | 67.0*<sub>a</sub> | 35.2 | 98.8 | 245.4*<sub>b</sub> | 122.2 | 368.5 |
| Scenario 6<br>Hydropower (Green Energy) and fisheries native salmonids | 226.4*<sub>a</sub> | 147.0 | 305.7 | 685.6*<sub>bd</sub> | 311.2 | 1060.0 | 284.3*<sub>ac</sub> | 179.2 | 389.3 | 699.6*<sub>d</sub> | 371.1 | 1028.1 |

Note. CI = confidence interval. CS estimates in each row that share subscripts do not differ significantly ( $p \geq .05$ ; Poe test).
<sup>a</sup>NOK and SEK were converted to € <sup>b</sup>For scenario descriptions see Table 3.
\* $p < .05$ .

Countries differed, however, regarding the general level of economic values generated through the scenarios. As evidenced by significant Poe-test results (Table 8), the Swedes benefited substantially more than the French and Norwegian citizens from all six scenarios while none of the CS differences between the latter two countries was significant (CS range in Sweden: 245.4 € to 699.6 € per year; in France: 77.6 € to 226.4 € per year; in Norway: 67.0 € vs. 303.7 € per year; Table 8). Compared to these consistent differences, welfare estimates in Germany varied from scenario to scenario relative to the other countries (CS range in Germany: 75.6 € to 685.6 € per year; Table 8). Despite these between-country differences in value levels, the significantly lower CS values found in every country for the hydropower (green-energy) Scenario 5, and their improvement when expanding the management goal by capture fisheries (Scenario 6), suggest uniform within-country variations of the combined impact of each scenario’s combination of attribute levels (Table 8).

## Discussion

We investigated the preferences of the general population of France, Germany, Norway and Sweden for hypothetical river development programs aimed at improving the ecological status of rivers in the vicinity of respondents’ places of residence, including riverine biodiversity and the rivers’ potential for fisheries. We found that citizens in all study countries had similar preferences for river attributes and generally benefited from these programs though at different levels of MWTP and CS. In all countries, MWTP estimates drove total river utility in the same direction resulting in similar patterns of differences in CS values across the six policy scenarios. Simultaneously, significant standard deviations associated with model coefficients signaled taste heterogeneity in the population. The development of nearby rivers was generally preferred to maintaining their perceived ecological status quo. The abiotic river attributes, particularly a minimized number of hydropower dams and good bathing water quality, contributed considerably to total river utility. Of the biotic attributes, only the fish species occurring in a river made a substantive, though country-specifically varying, contribution but not a species’ abundance or a river’s native biodiversity. Our hypotheses about people’s preference for good water quality, easy river bank access and native biodiversity thus received support as did our assumption of country-specific preferences for particular fish species. We propose a range of mechanisms for our findings below.

While five fish species added to a river’s perceived utility in France (rainbow trout, Atlantic salmon, brown trout, European eel, sturgeon) and Norway (brook trout, rainbow trout, Atlantic salmon, brown trout, sturgeon), more species than in the other countries, the Norwegians gained significantly more utility than the French from Atlantic salmon. This species’ high economic and cultural relevance in Norway is underscored by an MWTP value that is higher than those for most other species valued in this country (brook trout, rainbow trout, sturgeon). Across all countries, the native salmonid species (Atlantic salmon, brown trout) achieved seven nonzero MWTP values as opposed to four values for the nonnative salmonids (brook trout and rainbow trout), suggesting a cross-country preference for salmonid species that happen to be native (see Kochalski et al. 2018 for perceived nativeness of salmonid species). Because in Sweden and Germany fewer species than in France and Norway exhibited significant MWTP values (four and two, respectively), our results collectively give rise to speculations that the societal significance of fish species might be declining along a longitudinal gradient from European countries with extensive Atlantic seaboards (Norway, France) to more eastward countries with only indirect access to the Atlantic ocean but with long coastlines bordering the Baltic sea (Sweden, Germany), and with large areas remote from any seashore (Germany). An explanation for these between-country differences could be that European societies have been selectively affected by the “shifting baseline syndrom” (Pauly 1995). This process describes a long-term inter- and intragenerational extinction of knowledge of, and experience with, the conditions of the biological environments people live in due to a loss of opportunities to interact with nature (Papworth et al. 2009; Soga and Gaston 2016), including domestic fish species (Kochalski et al. 2018; Liebich et al. 2018) and possibly other components of river biodiversity. As a result, people may have become disconnected from (largely) extinct species like Atlantic salmon in countries such as Germany (Wolter 2015; Lenders et al. 2016; Kochalski et al. 2018; Liebich et al. 2018). Such a development would be critical as a loss of memory of past environmental degradation may ultimately lead to a reduction in the public’s engagement with, and notably their willingness-to-pay for, conservation efforts (McClenachan et al. 2018). While people were indifferent to grayling, European eel provided benefits to the French and Swedish societies as did sturgeon to all countries except Sweden. The latter two species either have been declining in recent years, such as eel in Germany, where it is economically still important to local-scale inland fisheries, or are threatened with extinction globally, like the sturgeon species (Freyhof and Brooks 2011). These findings suggest that expensive restoration activities tailored toward eel and sturgeon (e.g., BfN 2010; European Commission 2014) in countries where they were valued in our study are likely to receive considerable public support, whereas efforts to reintroduce, for instance, Atlantic salmon in Germany (e.g., Wolter 2015), the only country where this species was not valued, would not.

Confirming our hypothesis, native river biodiversity generated utility in three countries, a finding that was previously identified for rivers in Sweden (Kataria 2009) and in the UK (Hanley et al. 2006) as well as for the ocean (Jefferson et al. 2014; Jobstvogt et al. 2014; Daigle et al. 2016). The abundance of the focal fish species presented in the CE was only valued in Germany where it generated negative utility, suggesting a tendency among Germans to prefer a diversified assemblage of fish species that is not dominated by a single species, which is in line with a preference for general biodiversity. As both these attributes made only small contributions to CS values and in opposing directions, biodiversity in general provided little welfare to the four societies. People even benefited from nonnative fish species, whose occurrence may threaten freshwater biodiversity (Gozlan et al. 2010; Cucherousset and Olden 2011. The utility attached to these species may originate from the fact that they have been stocked for a long period of time (Aas et al. in press) and are likely considered naturalized across Europe (Kochalski et al. 2018), and also from their relevance in the diets of many Europeans. Though previous studies have shown the importance of perceived biodiversity to human health and well-being, this relationship is not direct and depends on the presence of particular species (Fuller et al. 2007; Pett et al. 2016). Moreover, the perceptions of biodiversity are often at odds with the actual biodiversity present in an ecosystem (Fuller et al. 2007; Shwartz et al. 2014; Belaire et al. 2015; Sandifer et al. 2015).

According to our results, the bathing water quality and the presence of hydropower plants as abiotic river characteristics were more important to people than ecological properties like biodiversity. Bathing water quality drove preferences significantly, particularly in Sweden and Germany, confirming results from previous studies in both freshwater and marine waters (Daigle et al. 2016; Artell and Huhtala 2017). A usability-based index of water quality like the one we employed was previously found to be correlated with an indicator that measured the ecological status according to the WFD (Artell and Huhtala 2017), suggesting that management towards good water quality may indirectly elevate biodiversity. The societal value attached to bathing water quality is probably linked to its perceived relationship with what people consider a clean and healthy river (Jefferson et al. 2014; Daigle et al. 2016; Liebich et al. 2018). The usage of rivers for hydropower production also contributed significantly, though negatively, to a river’s total utility, most of all in Germany and least so in Norway although the Norwegian respondents reported the comparatively strongest perceived impact (status quo) of hydropower dams on their rivers’ water flow. While recently hydropower has had its rebirth in an attempt to increase renewable energy production worldwide (Zarfl et al. 2015; Winemiller et al. 2016; Couto and Olden 2018), in our survey, a river’s utility decreased substantially with increasing fragmentation due to hydropower dams. Given the major negative impact that river fragmentation has had on riverine biodiversity over centuries (Wolter 2015; Lawrence et al. 2016; Lenders et al. 2016), and is continuing to have globally (Zarfl et al. 2015; Winemiller et al. 2016; Cooper et al. 2017; Couto and Olden 2018), it is vital from an environmental perspective to implement measures (e.g., installing fish ladders for migratory fishes or applying administrative means to adjust a river’s flow regime; Poff and Schmidt 2016) to better balance social and ecological requirements as was done in Norwegian watersheds (Alfredsen et al. 2012; Ruud and Fjeldstad 2015; Norwegian Environment Agency 2017). While such regulations are not in place in the small-scale hydropower operations common to, for example, Germany (Zarfl et al. 2015), the very low negative MWTP of hydropower in Norway may have resulted from this country’s strong dependence on, and hence from a broad societal acceptance of, hydropower, or from the fact that Norway has protected more watersheds against hydropower development than, for instance, Sweden (Norwegian Environment Agency 2017; Swedish Environment Law 2017). Supporting our findings, Kataria (2009) found that Swedish households were willing to pay for environmental improvements in hydropower-regulated waters (like, e.g., ecologically optimized river vegetation or increased biodiversity). With the exception of Germany, easy access to the river banks was also preferred, particularly in Sweden. This result agreed with previous findings from Poland (Birol et al., 2009). Because we informed our respondents in the survey that easy access to the river banks may cause the loss of natural habitats of riparian plants and animals (due to artificial embankments or trampling), the preferences found in this study imply that respondents also valued river conditions that are at least partly anthropocentric in nature (by providing roads to and pathways along the river banks) and may thus exert pressure on ecological river functioning.

Complementing the attribute-based utilities just discussed, results from the scenario analyses showed that the preferences for the outcomes of selected river basin management plans were very similar in the populations of the four European countries albeit at different CS levels. Our findings showcase the benefits that ecological river management and restoration can provide to the four countries, most strongly in Sweden but also in Germany. The findings also demonstrate how a management plan that fails to meet the general population’s preferences (as in the case of the green-energy hydropower Scenario 5) can cause a significant decline of a river’s perceived economic value (compared to the native-salmonid fisheries Scenario 1 and to both conservation-oriented Scenarios 3 and 4). But the results also reveal how the CS values can be improved when the management strategy is revised to also supply ecosystem services such as capture fisheries for native salmonids (Scenario 6).

In terms of the limitations of our study, the low importance found for the biotic attributes needs to be interpreted with some caution. Although previous research has found that the general public was indeed able to derive economic values from unfamiliar ecological objects such as biodiversity (e.g., Börger and Hattam 2017), the potential of a CE to inform respondents about the ecological relevance of river attributes is limited. Despite our attempt to familiarize respondents with the attributes beforehand, respondents are unlikely to have been fully knowledgeable about biodiversity and the benefits it provides to society. In consequence, respondents may also have expressed their preferences for biodiversity indirectly through choosing high levels of water quality and fewer dams assuming that biodiversity would benefit from both. Moreover, the study context may have overemphasized negative aspects of hydropower production. We introduced the questionnaire as a survey on “humans-rivers-species diversity” which may have biased respondents’ answers in a proecological direction. Had the CE been administered in a survey on, for example, electricity production, hydropower may have performed more positively in relation to fossil fuel or nuclear power.

## Implications and conclusions

Our findings have five main implications. First, all significant MWTP values had the same algebraic signs in all countries. These unidirectional impacts suggest a common preference structure, which was corroborated by a uniform pattern of within-country variations of the CS values across the scenarios. Second, the differing levels of the utility estimates between the countries likely result from cultural and biogeographical differences between the countries emphasizing the necessity to consider the specific societal conditions in each country in the public discourse about the values of nature and biodiversity conservation. Third, citizens in all countries preferred ecologically valuable conditions, which simultaneously supply ecosystem services (e.g., good bathing water quality, fisheries), with the preference for easy bank access and for nonnative salmonids being important exceptions. Forth, the relevance of selected fish species varied between the four countries, which has implications for the acceptability of species-centered conservation efforts. Lastly, the scenario analyses demonstrated that our CE data allow for a comparison of a range of alternative river basin management goals. These data can thus be used for informing policy makers’ decisions on improvements of the ecological status of domestic rivers while gauging each decision’s social benefit (or cost).

To conclude, our results show that ecological river management can create high levels of economic benefits in particular through an optimal combination of the three abiotic river attributes (hydropower dams, bank accessibility, bathing water quality) but also through efforts to restore declining or extinct fish species. Thus, if environmental managers also considered biotic river characteristics, including the fish species assemblage, even more benefits could be generated. As the common cross-country utility structure allows for taking advantage of synergy effects in planning efforts within the European Union and cooperating countries (Norway), our results give indications of which policies are likely to generate high societal benefits that might justify even expensive river restoration efforts.

## Complicance with Ethical Standards

The authors declare that they have no conflict of interest. All procedures in this study involving human participants were conducted according to the ethical standards of the German Research Foundation (DFG) and in compliance with national data protection acts.

## Acknowledgments

This study was funded by the German Research Foundation (DFG; grant to R.A., number AR 712/4-1) within the project SalmoInvade in the BiodivERsA 2012-2013 Pan-European call (supported by the EU’s Horizon 2020 research and innovation program). R.A. also received funding from the German Federal Ministry of Education and Research (BMBF) within the project Besatzfisch (grant number 01UU0907) in the Programme for Social Ecological Research and through the EU’s Horizon 2020 program under the Marie Sklodowska-Curie grant agreement (No 642893). We are grateful to Ulf Liebe, Julian Sagebiel, Wolfgang Bandilla, Michael Braun and to our colleagues at Leibniz-Institute of Freshwater Ecology and Inland Fisheries and Humboldt-Universität zu Berlin for insightful comments on the study concept and data interpretation. We would also like to thank Dorothée Behr, Julien Cucherousset, Jörgen Johnsson, Kjetil Hindar, all other members of the SalmoInvade project and the team of Language Connect for assisting with the translation of the questionnaire and for discussing its content. Special thanks go to Frederik Funke, Marco Reich, Alexandra Wachenfeld and all others at LINK, forsa and Norstat for collecting the data and to all study participants for their kind cooperation.

## References

Aas Ø, Cucherousset J, Fleming IA, et al (in press) Salmonid stocking in five North Atlantic jurisdictions - identifying drivers and barriers to policy change. Aquat Conserv.

ADM Arbeitskreis Deutscher Markt- und Sozialforschungsinstitute e. V. (2018) The ADM-Sampling-System for Telephone Surveys. https://www.adm-ev.de/telefonbefragungen/?L=1. Accessed 30 May 2018

Ahtiainen H, Pouta E, Artell J (2015) Modelling asymmetric preferences for water quality in choice experiments with individual-specific status quo alternatives. Water Resour Econ 12:1–13. doi: 10.1016/j.wre.2015.10.003

Alfredsen K, Harby A, Linnansaari T, Ugedal O (2012) Development of an inflow-controlled environmental flow regime for a Norwegian river. River Res Appl 28:731–739. doi: 10.1002/rra.1550

Arlinghaus R, Engelhardt C, Sukhodolov A, Wolter C (2002) Fish recruitment in a canal with intensive navigation: implications for ecosystem management. J Fish Biol 61:1386–1402. doi: doi:10.1006/jfbi.2002.2148

Artell J, Huhtala A (2017) What are the benefits of the Water Framework Directive? Lessons learned for policy design from preference revelation. Environ Resour Econ 68:847–873. doi: 10.1007/s10640-016-0049-8

Auerbach DA, Deisenroth DB, McShane RR, et al (2014) Beyond the concrete: accounting for ecosystem services from free-flowing rivers. Ecosyst Serv 10:1–5. doi: 10.1016/j.ecoser.2014.07.005

Belaire JA, Westphal LM, Whelan CJ, Minor ES (2015) Urban residents’ perceptions of birds in the neighborhood: biodiversity, cultural ecosystem services, and disservices. Condor 117:192–202. doi:10.1650/CONDOR-14-128.1

BfN Bundesamt für Naturschutz (ed) (2010) German action plan for the conservation and restoration of the European sturgeon (Acipenser sturio). Federal Ministry for the Environment, Nature Conservation and Nuclear Safety (BMU), Bonn, Germany

Birol E, Hanley N, Koundouri P, Kountouris Y (2009) Optimal management of wetlands: quantifying trade-offs between flood risks, recreation, and biodiversity conservation. Water Resour Res 45:. doi:10.1029/2008WR006955

Bliemer MCJ, Rose JM (2013) Confidence intervals of willingness-to-pay for random coefficient logit models. Transp Res Part B 58:199–214. doi: 10.1016/j.trb.2013.09.010

Börger T, Hattam C (2017) Motivations matter: behavioural determinants of preferences for remote and unfamiliar environmental goods. Ecol Econ 131:64–74. doi: 10.1016/j.ecolecon.2016.08.021

Brouwer R (2008) The potential role of stated preference methods in the Water Framework Directive to assess disproportionate costs. J Environ Plan Manag 51:597–614. doi: 10.1080/09640560802207860

Collen B, Whitton F, Dyer EE, et al (2014) Global patterns of freshwater species diversity, threat and endemism. Glob Ecol Biogeogr 23:40–51. doi: 10.1111/geb.12096

Cooper AR, Infante DM, Daniel WM, et al (2017) Assessment of dam effects on streams and fish assemblages of the conterminous USA. Sci Total Environ 586:879–889. doi: 10.1016/j.scitotenv.2017.02.067

Couto TBA, Olden JD (2018) Global proliferation of small hydropower plants - science and policy. Front Ecol Environ 16:91–100. doi: 10.1002/fee.1746

Cucherousset J, Olden JD (2011) Ecological impacts of non-native freshwater fishes. Fisheries 36:215–230. doi: 10.1080/03632415.2011.574578

Daigle RM, Haider W, Fernández-Lozada S, et al (2016) From coast to coast: public perception of ocean-derived benefits in Canada. Mar Policy 74:77–84. doi: 10.1016/j.marpol.2016.09.012

de Jager AL, Vogt J V. (2010) Development and demonstration of a structured hydrological feature coding system for Europe. Hydrol Sci J 55:661–675. doi: 10.1080/02626667.2010.490786

Dillman DA, Smyth JD, Christian LM (2014) Internet, Phone, Mail, and Mixed-Mode Surveys. Wiley & Sons, Hoboken, New Jersey, USA

Dinter J, Bechthold A, Boeing H, et al (2016) Fish intake and prevention of selected nutrition-related diseases. Ernährungs Umschau 63:148–154. doi: 10.4455/eu.2016.032

Dudgeon D, Arthington AH, Gessner MO, et al (2006) Freshwater biodiversity: importance, threats, status and conservation challenges. Biol Rev Camb Philos Soc 81:163–182. doi: 10.1017/S1464793105006950

European Commission (2000) Directive 2000/60/EC of the European Parliament and of the Council establishing a framework for the Community action in the field of water policy. http://eur-lex.europa.eu/legal-content/EN/TXT/PDF/?uri=CELEX:32000L0060&from=EN. Accessed 31 Oct 2016

European Commission (2014) Report from the commission to the Council and the European parliament on the outcome of the implementation of the Eel Management Plan. https://eur-lex.europa.eu/resource.html?uri=cellar:d77e3ffd-5918-11e4-a0cb-01aa75ed71a1.0006.03/DOC_1&format=PDF. Accessed 25 Jul 2018

European Commission (2017) WFD: Timetable for implementation. http://ec.europa.eu/environment/water/water-framework/info/timetable_en.htm. Accessed 9 Jan 2017

European Environment Agency (2018) European waters - assessment of status and pressures. Publications Office of the European Union, Luxembourg, Luxembourg

Eurostat (2015) Census data for the online populations. http://ec.europa.eu/eurostat/de/data/database. Accessed 30 May 2018

Eurostat (2016) Information on internet penetration. http://ec.europa.eu/eurostat/statistics-explained/index.php/Digital_economy_and_society_statistics_-_households_and_individuals. Accessed 1 Jun 2018

Freyhof J, Brooks E (2011) European red list of freshwater fishes. Publications Office of the European Union, Luxembourg

Fuller RA, Irvine KN, Devine-Wright P, et al (2007) Psychological benefits of greenspace increase with biodiversity. Biol Lett 3:390–394. doi: 10.1098/rsbl.2007.0149

Gozlan RE, Britton JR, Cowx I, Copp GH (2010) Current knowledge on non-native freshwater fish introductions. J Fish Biol 76:751–786. doi: 10.1111/j.1095-8649.2010.02566.x

Hanemann WM (1984) Welfare evaluations in contingent valuation experiments with discrete responses. Am J Agric Econ 66:332–341

Hanley N, Wright RE, Alvarez-Farizo B (2006) Estimating the economic value of improvements in river ecology using choice experiments: an application to the water framework directive. J Environ Manage 78:183–193. doi: 10.1016/j.jenvman.2005.05.001

Heckel C, Glemser A, Meier G (2014) Das ADM-Telefonstichproben-System. In: ADM Arbeitskreis Deutscher Markt- und Sozialforschungsinstitute e. V. (ed) Stichproben-Verfahren in der Umfrageforschung. Springer, Wiesbaden, Germany, pp 137–166

Hering D, Borja A, Carstensen J, et al (2010) The European Water Framework Directive at the age of 10: a critical review of the achievements with recommendations for the future. Sci Total Environ 408:4007–4019. doi: 10.1016/j.scitotenv.2010.05.031

Hindar K (2003) Wild Atlantic salmon in Europe: status and perspectives. In: Gallaugher P, Wood L (eds) The World Summit on Salmon. Continuing Studies in Science, Burnaby, British Columbia, Canada, pp 47–52

Holmlund CM, Hammer M (1999) Ecosystem services generated by fish populations. Ecol Econ 29:253–268

Huet M (1949) Aperçu des relations entre la pente et les populations piscicoles des eaux courantes. Schweizerische Zeitschrift für Hydrol 11:332–351. doi: 10.1007/BF02503356

Jefferson RL, Bailey I, Laffoley D d’A, et al (2014) Public perceptions of the UK marine environment. Mar Policy 43:327–337. doi: 10.1016/j.marpol.2013.07.004

Jobstvogt N, Hanley N, Hynes S, et al (2014) Twenty thousand sterling under the sea: estimating the value of protecting deep-sea biodiversity. Ecol Econ 97:10–19. doi: 10.1016/j.ecolecon.2013.10.019

Kalinkat G, Cabral JS, Darwall W, et al (2017) Flagship umbrella species needed for the conservation of overlooked aquatic biodiversity. Conserv Biol 31:481–485. doi: 10.1111/cobi. 12813

Kataria M (2009) Willingness to pay for environmental improvements in hydropower regulated rivers. Energy Econ 31:69–76. doi: 10.1016/j.eneco.2008.07.005

Kochalski S, Riepe C, Fujitani M, et al (2018) Public perception of river fish biodiversity in four European countries. Conserv Biol. doi: 10.1111/cobi.13180

Lawrence ER, Kuparinen A, Hutchings JA (2016) Influence of dams on population persistence in Atlantic salmon (Salmo salar). Can J Zool 94:329–338

Lenders HJR, Chamuleau TPM, Hendriks AJ, et al (2016) Historical rise of waterpower initiated the collapse of salmon stocks. Sci Rep 6:. doi: 10.1038/srep29269

Liebich KB, Kocik JF, Taylor WW (2018) Reclaiming a space for diadromous fish in public psyche and sense of place. Fisheries 43:231–240. doi: https://doi.org/10.1002/fsh.10063

Liu P, Lien K, Asche F (2016) The impact of media coverage and demographics on the demand for Norwegian salmon. Aquac Econ Manag 20:342–356. doi: 10.1080/13657305.2016.1212126

Louviere JJ, Hensher DA, Swait JD (2000) Stated choice methods. Cambridge University Press, Cambridge, UK

Marschak J (1960) Binary choice constraints on random utility indicators. In: Arrow KJ, Karlin S, Suppes P (eds) Stanford Symposium on Mathematical Methods in the Social Sciences. Stanford University Press, Stanford, California, USA, pp 312–329

McClenachan L, Matsuura R, Shah P, Dissanayake STM (2018) Shifted baselines reduce willingness to pay for conservation. Front Mar Sci 5:. doi: 10.3389/fmars.2018.00048

McFadden D (1974) Conditional logit analysis of qualitative choice behavior. In: Zarembka P (ed) Frontiers in econometrics. Academic Press, New York, New York, USA, pp 105–142

Meyerhoff J, Boeri M, Hartje V (2014) The value of water quality improvements in the region Berlin-Brandenburg as a function of distance and state residency. Water Resour Econ 5:49–66. doi: http://dx.doi.org/10.1016/j.wre.2014.02.001

Neteler M, Bowman MH, Landa M, Metz M (2012) GRASS GIS: A multi-purpose open source GIS. Environ Model Softw 31:124–130. doi: 10.1016/j.envsoft.2011.11.014

Nichols WJ (2014) Blue mind. Back Bay Books, New York, New York, USA

Nieminen E, Hyytiäinen K, Lindroos M (2016) Economic and policy considerations regarding hydropower and migratory fish. Fish Fish. doi: 10.1111/faf.12167

Norwegian Environment Agency (2017) Protection plan for water resources. http://www.miljodirektoratet.no/en/Areas-of-activity1/Inland-waters/Protection-Plan-for-Water-Resources/. Accessed 20 Jan 2017

Olsen SB, Meyerhoff J (2016) Will the alphabet soup of design criteria affect discrete choice experiment results? Eur Rev Agric Econ. doi: 10.1093/erae/jbw014

Papworth SK, Rist J, Coad L, Milner-Gulland EJ (2009) Evidence for shifting baseline syndrome in conservation. Conserv Lett 2:93–100. doi: 10.1111/j.1755-263X.2009.00049.x

Pauly D (1995) Anecdotes and the shifting baseline syndrome of fisheries. Trends Ecol Evol 10:430

Pett TJ, Shwartz A, Irvine KN, et al (2016) Unpacking the people-biodiversity paradox: a conceptual framework. Bioscience 66:576–583. doi: 10.1093/biosci/biw036

Poe GL, Giraud KL, Loomis JB (2005) Computational methods for measuring the difference of empirical distributions. Am J Agric Econ 87:353–365

Poff NL, Schmidt JC (2016) How dams can go with the flow. Science (80-) 353:1099–1100. doi: 10.1126/science.aah4926

Polizzi C, Simonetto M, Barausse A, et al (2015) Is ecosystem restoration worth the effort? The rehabilitation of a Finnish river affects recreational ecosystem services. Ecosyst Serv 14:158–169. doi: 10.1016/j.ecoser.2015.01.001

Rockström J, Steffen W, Noone K, et al (2009) A safe operating space for humanity. Nature 461:472–475

Ruud AA, Fjeldstad H-P (2015) Vannforskriften og norsk vannkraftproduksjon. Kan miljedesign og funksjonsmâl gi bedre planprosesser? VANN 152–162. doi: 10.13140/RG.2.1.3531.8881

Sagebiel J, Glenk K, Meyerhoff J (2017) Spatially explicit demand for afforestation. For Policy Econ 78:190–199. doi: 10.1016/j.forpol.2017.01.021

Sandifer PA, Sutton-Grier AE, Ward BP (2015) Exploring connections among nature, biodiversity, ecosystem services, and human health and well-being: opportunities to enhance health and biodiversity conservation. Ecosyst Serv 12:1–15. doi: 10.1016/j.ecoser.2014.12.007

Scarpa R, Rose J (2008) Design efficiency for non-market valuation with choice modelling: how to measure it, what to report and why. Aust J Agric Resour Econ 253–282. doi: 10.1111/j.1467-8489.2007.00436.x

Shwartz A, Turbé A, Simon L, Julliard R (2014) Enhancing urban biodiversity and its influence on city-dwellers: an experiment. Biol Conserv 171:82–90. doi: 10.1016/j.biocon.2014.01.009

Soga M, Gaston KJ (2016) Extinction of experience: the loss of human-nature interactions. Front Ecol Environ 14:94–101. doi: 10.1002/fee.1225

Swedish Environment Law Miljöbalk (1998:808) (2017) http://www.riksdagen.se/sv/dokumentlagar/dokument/svensk-forfattningssamling/miljobalk-1998808_sfs-1998-808. Accessed 20 Jan 2017

Szałkiewicz E, Jusik S, Grygoruk M (2018) Status of and perspectives on river restoration in Europe: 310,000 Euros per hectare of restored river. Sustain 10:. doi: 10.3390/su10010129

Tockner K, Pusch M, Borchardt D, Lorang MS (2010) Multiple stressors in coupled river-floodplain ecosystems. Freshw Biol 55:135–151. doi: 10.1111/j.1365-2427.2009.02371.x

Train K (2009) Discrete choice methods with simulation. Cambridge University Press, Cambridge, UK

UNESCO (2016) ISCED: International standard classification of education. http://www.uis.unesco.org/Education/Pages/international-standard-classification-of-education.aspx. Accessed 31 Oct 2016

White M, Smith A, Humphryes K, et al (2010) Blue space: the importance of water for preference, affect, and restorativeness ratings of natural and built scenes. J Environ Psychol 30:482–493. doi: 10.1016/j.jenvp.2010.04.004

Winemiller KO, McIntyre PB, Castello L, et al (2016) Balancing hydropower and biodiversity in the Amazon, Congo, and Mekong. Science (80-) 351:128–129

Wolter C (2015) Historic catches, abundance, and decline of Atlantic salmon Salmo salar in the River Elbe. Aquat Sci 77:367–380. doi: 10.1007/s00027-014-0372-5

WWF World Wide Fund for Nature (2001) The status of wild Atlantic salmon. http://d2ouvy59p0dg6k.cloudfront.net/downloads/salmon2.pdf%0A. Accessed 7 Dec 2016

Zarfl C, Lumsdon AE, Berlekamp J, et al (2015) A global boom in hydropower dam construction. Aquat Sci 77:161–170. doi: 10.1007/s00027-014-0377-0

